# Chromatin environment, transcriptional regulation, and splicing distinguish lincRNAs and mRNAs

**DOI:** 10.1101/088484

**Authors:** Marta Melé, Kaia Mattioli, William Mallard, David M Shechner, Chiara Gerhardinger, John Rinn

## Abstract

While long intergenic noncoding RNAs (lincRNAs) and mRNAs share similar biogenesis pathways, these transcript classes differ in many regards. LincRNAs are less evolutionarily conserved, less abundant, and more tissue-specific, suggesting that their pre‐ and post-transcriptional regulation is different from that of mRNAs. Here, we perform an in-depth characterization of the features that contribute to lincRNA regulation in multiple human cell lines. We find that lincRNA promoters are depleted of transcription factor (TF) binding sites, yet enriched for some specific factors such as GATA and FOS relative to mRNA promoters. Surprisingly, we find that H3K9me3—a histone modification typically associated with transcriptional repression—is more enriched at the promoters of active lincRNA loci than at those of active mRNAs. Moreover, H3K9me3-marked lincRNA genes are more tissue-specific. The most discriminant differences between lincRNAs and mRNAs involve splicing. LincRNAs are less efficiently spliced, which cannot be explained by differences in U1 binding or the density of exonic splicing enhancers, but may be partially attributed to lower U2AF65 binding and weaker splicing–related motifs. Conversely, the stability of lincRNAs and mRNAs is similar, differing only with regard to the location of stabilizing protein binding sites. Finally, we find that certain transcriptional properties are correlated with higher evolutionary conservation in both DNA and RNA motifs, and are enriched in lincRNAs that have been functionally characterized.

## INTRODUCTION

Transcription and post-transcriptional regulation are crucial processes for the biogenesis and function of all RNA species. While several non-coding RNA classes— such as microRNAs, snoRNAs, and tRNAs—have post-transcriptional processing pathways that are unique to each class, long intergenic noncoding RNA (lincRNA) biogenesis shares many similarities with that of protein-coding mRNAs (Quinn and Chang 2015). LincRNAs are transcribed by RNA Polymerase II, 5’-capped, spliced, and polyadenylated. Like those of mRNAs, lincRNA splice site dinucleotides are canonical, suggesting that they use the same splicing regulatory signals as mRNAs (Derrien et al. 2012). However, lincRNAs differ from mRNAs in several regards: they have fewer (although usually longer) exons (Derrien et al. 2012), they exhibit more tissue-specific expression (Ulitsky and Bartel 2013; Cabili et al. 2011) and they have higher nuclear localization than their mRNA counterparts (Djebali et al. 2012). Whether such observations are a reflection of differences in the transcriptional processing of lincRNAs, and whether differential transcriptional regulation may have functional implications, remains unknown.

At the primary sequence level, most lincRNAs are less evolutionarily conserved than mRNAs (Hezroni et al. 2015; Guttman et al. 2009), leading some to argue that lincRNAs may be functionless byproducts of transcriptional noise (Kowalczyk et al. 2012). However, recent work has found that a subset of lincRNAs contain conserved regulatory elements, including transcription factor binding sites (Necsulea et al. 2014), nuclear localization signals (Hacisuleyman et al. 2016), and splicing motifs (Ponjavic et al. 2007; Haerty and Ponting 2015). This suggests that the pre‐ and post-transcriptional regulation of some lincRNAs is functionally relevant. Despite this, a global assessment of the importance of lincRNA pre‐ and post-transcriptional regulation, and how such regulation compares to that of mRNAs, remains unresolved.

Here, we combine a battery of functional genomic analysis and biochemical assays to comprehensively interrogate the differences between the lifecycles of lincRNAs and mRNAs in humans. Our analysis provides an extensive characterization of the lincRNA lifecycle and their distinguishing properties from mRNAs.

## RESULTS

### Active lincRNA promoters are depleted for most histone marks, but are enriched in H3K9me3

To systematically survey the properties associated with the RNA lifecycle, and to avoid potentially confounding influences arising from the regulation of overlapping genes, we focused our analysis on intergenic lncRNAs (termed “lincRNAs”). These lincRNAs were compared to a reference set of mRNAs (Supplemental Methods), resulting in a dataset comprising 5,196 lincRNAs and 19,575 mRNAs. For most analyses, we only compared lincRNAs to mRNAs with similar expression levels. We first addressed whether the promoters of transcribed lincRNAs (defined as “active promoters”) and a set of expression-matched mRNAs differed in their chromatin environment and transcriptional regulation. We defined promoter regions as 5Kb upstream and downstream (+/− 5Kb) from the TSS. We then curated chromatin immunoprecipitation (ChIP) experiments for 70 histone modifications and 370 transcription factors, in seven and eleven ENCODE cell lines, respectively (The ENCODE Project Consortium 2004). Finally, we used RNA-seq from 20 human tissues to calculate tissue–specificity (Cabili et al. 2011).

We found that, in each of the seven tested cell lines and relative to their mRNA counterparts, the promoters of most lincRNAs were depleted for nearly all histone marks (Figure 1A; Supplemental Fig S1). Intriguingly, the only histone mark enriched in active lincRNA promoters compared to mRNAs was H3K9me3, a modification commonly found in transcriptionally repressed regions (Figure 1A, B, Supplemental Fig S2, Supplemental Table S1). Despite this, lincRNA genes with and without H3K9me3 were expressed at similar levels in all but one cell line (Wilcoxon test; FDR<0.05; Supplemental Fig S3). Interestingly, H3K9me3-marked lincRNAs were significantly more tissue-specific than those lacking this histone mark, in five of the seven cell lines (FDR<0.05; Supplemental Fig S4). In expression-matched mRNAs, neither expression levels nor tissue specificity differed significantly with regard to H3K9me3 status (Wilcoxon test; FDR>0.05; Supplemental Fig S3-S4).

**Figure 1.**
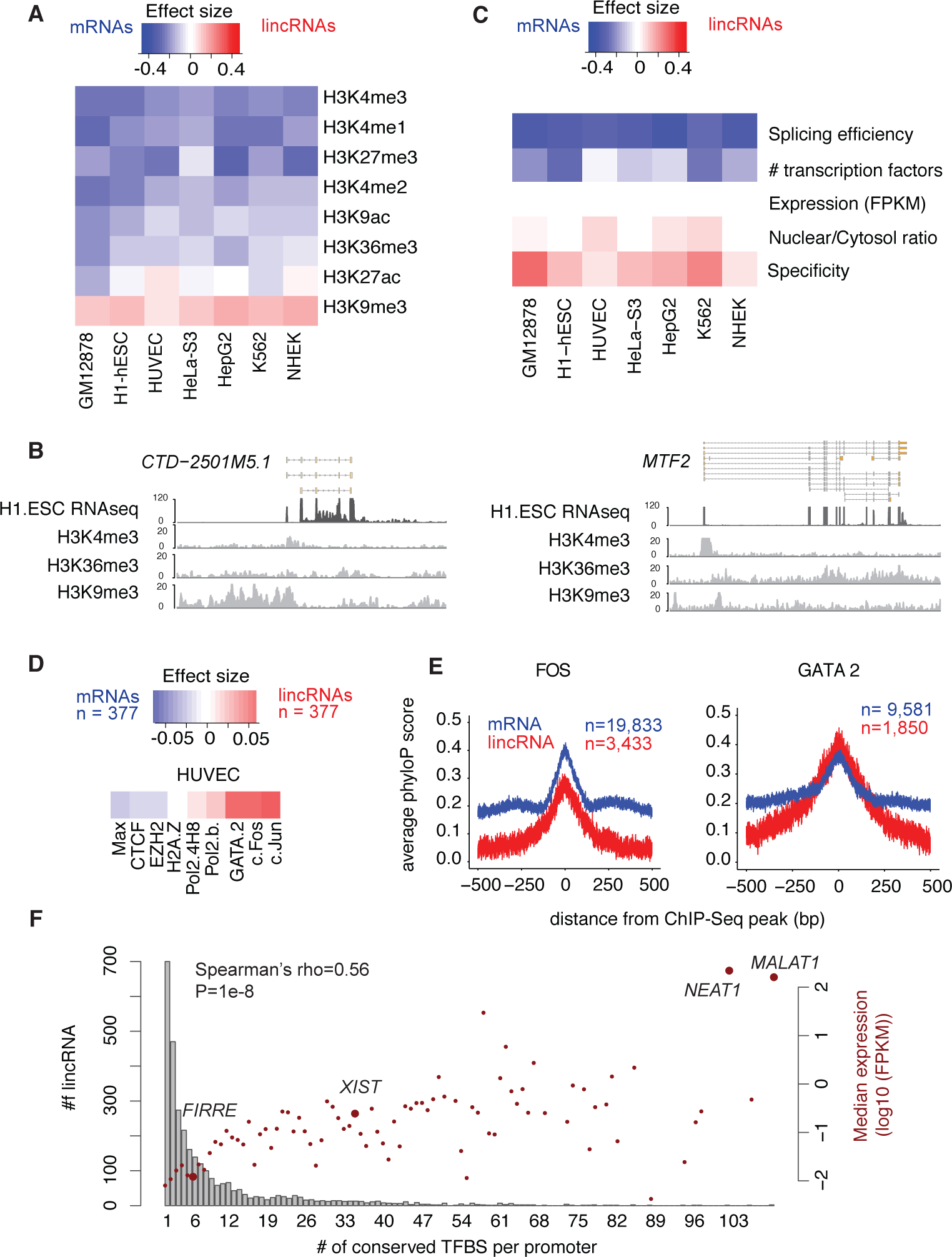
Certain histone marks and TFs are enriched in lincRNA promoters. **A.** Fisher effect size differences, comparing the presence or absence of eight histone marks (observed via ChIP) in the promoters of lincRNAs and control mRNAs, in seven ENCODE cell lines. Blue corresponds to larger values in mRNAs and red to larger values in lincRNAs. The number of genes ranged from 754 (HUVEC) to 1262 (GM12878) and corresponds to all lincRNAs expressed at more than 0.1 FPKM in the tested cell line, and to expression-matched mRNAs. **B.** ChIP-seq read coverage for three histone marks in one lincRNA (*left*) and one mRNA (*right*) with similar expression levels in H1 ESCs. **C.** Wilcoxon effect size differences between lincRNAs and mRNAs across several gene/promoter properties. The genes analyzed are the same as in A. **D.** Fisher effect size differences between lincRNA and mRNA promoters comparing the presence or absence of TFs, in HUVEC cells. **E.** Average conservation at FOS (*left*) and GATA2 (*right*) binding sites overlapping lincRNA and mRNA promoters centered on ChIP-seq peaks. Width of the lines represents the standard error. **F.** Histogram of the number of lincRNAs with different numbers of TFs with conserved binding sites in their promoters (bars, left axis) and the corresponding median expression for that group of lincRNAs (dots, right axis). The identity of notable functional lincRNAs within each group is highlighted.

### LincRNA promoters are enriched for specific transcription factors and have conserved binding sites

Using ChIP data, we next examined the binding of transcription factors at lincRNA promoters. In general, lincRNA promoters were bound by fewer classes of TFs than those of expression-matched mRNAs (Figure 1C, Supplemental Table S2). However, some TFs—including the GATA family, JUN, and FOS—were consistently enriched at lincRNA promoters in all cell lines (Supplemental Table S3, Figure 1D). To assess the functional relevance of these interactions, we calculated average conservation scores across TF binding sites (TFBSs) that intersected either lincRNA or mRNA promoters. We found that in lincRNA promoters, 87.4% of TFs had binding sites that were conserved compared to their flanking regions (FDR<0.05) whereas in mRNA promoters 97.8% of TFs were conserved (Supplemental Fig S5; Supplemental Table S4, S5). Nevertheless, in some cases, such as for GATA2, KAP1, and MBD4, the average conservation was higher in lincRNA promoters than in mRNA promoters (Figure 1E, Supplemental Fig S5). These results were consistent irrespective of how promoters were defined (i.e., in windows of 10 kb or windows of 3 kb) or whether TFBSs were based on ChipSeq peaks or on known TF binding motifs (Supplemental Fig S6, S7).

We next analyzed the conservation of TFBSs in lincRNA promoters, testing whether nucleotides within each TFBS were more conserved than those in the surrounding region (FDR <0.05). We observed that 61.9% of lincRNA promoters had at least one conserved TFBS (median of 1, mean of 6.2 conserved TFBS per lincRNA promoter. Figure 1F; Supplemental Table S6, S7). Similarly, 65.9% of the promoters of expression-matched mRNAs had at least one conserved TFBS (median of 2, mean of 8.55 conserved TFBS per promoter; Supplemental Fig S8). These proportions were larger than those found in random intergenic regions (Supplemental Table S8). Furthermore, expression of both lincRNAs and mRNAs increased with the number of conserved TFBS (Figure 1F; Supplemental Fig S8). Conversely, tissue specificity decreased with increasing number of conserved TFBS types (Supplemental Fig S9-S10). Finally, we found that lincRNAs present in lncRNAdb (a database of functionally characterized lincRNAs, (Amaral et al. 2011)) had significantly more conserved TFBSs than lincRNAs lacking functional characterization (Wilcoxon P=1.09e-05). Together, these results suggest that the number of conserved promoter TFBSs might be a useful characteristic with which to identify functional lincRNAs.

### LincRNA splicing is inefficient

We next sought to systematically investigate splicing efficiency in lincRNAs, which has been previously examined in smaller scale studies (Tilgner et al. 2012; Seidl et al. 2006). To this end, we developed a gene-level metric for calculating splicing efficiency (Supplemental Fig S11A). Reasoning that differences in splicing might be more evident in the nuclear pool of RNA, we first analyzed fractionated nuclear RNA-seq data from seven ENCODE cell lines (Djebali et al. 2012). Compared to mRNAs with similar expression levels, lincRNAs generally exhibited inefficient splicing in all cell lines (Figure 2A; Supplemental Table S1, Supplemental Fig S12A, S13).

**Figure 2.**
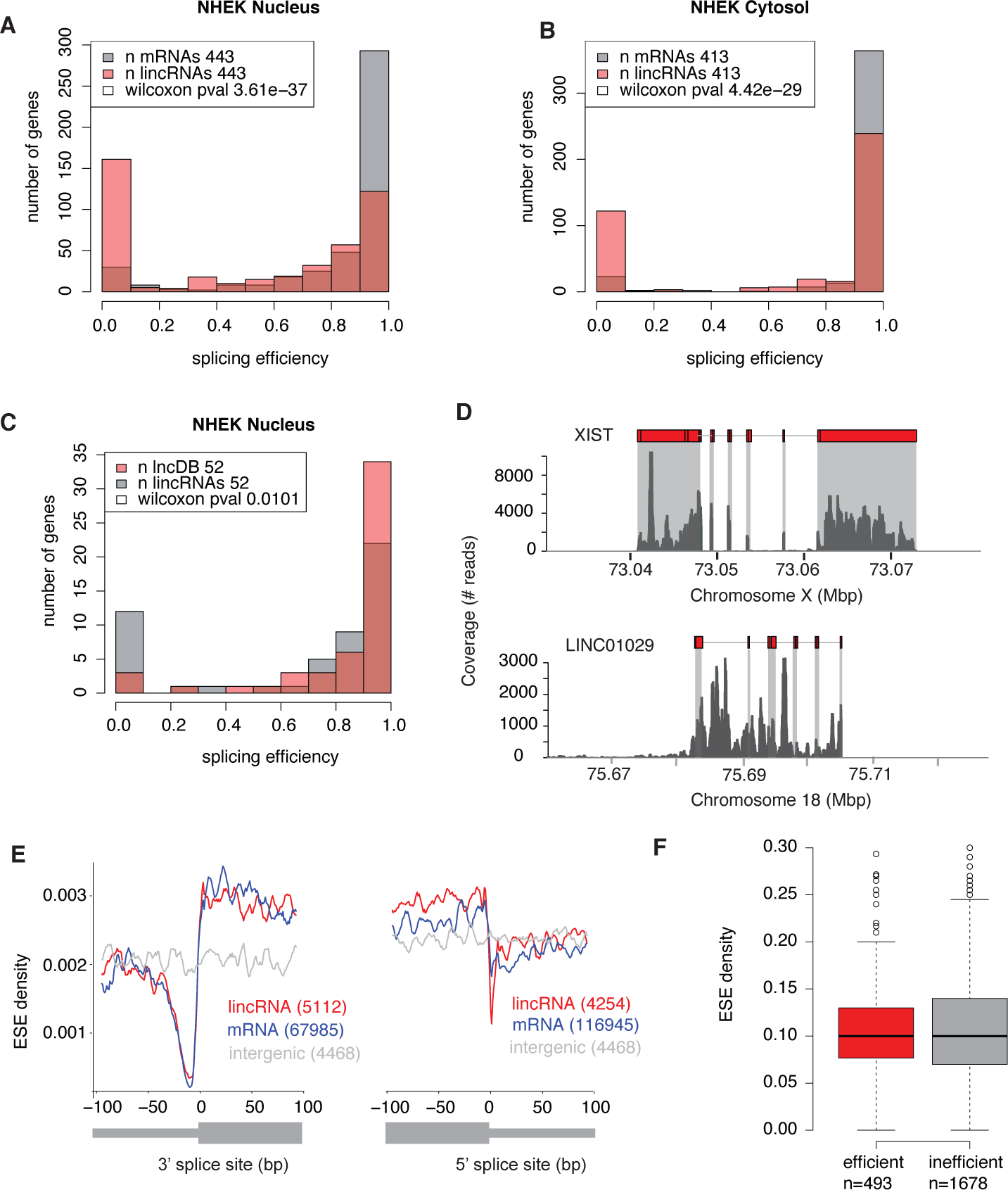
lincRNAs are less efficiency spliced than are mRNAs. **A.-B.** Splicing efficiency in lincRNAs (>0.1 FPKM) and expression-matched mRNAs in the nuclear fraction (A) and cytosolic fraction (B) of human NHEK cells. **C.** Splicing efficiency in functionally characterized (present in lncRNAdb) lincRNAs and expression-matched (>0.1 FPKM) uncharacterized lincRNAs. **D.** RNA-seq read coverage for a representative functional, efficiently-spliced lincRNA, *XIST* (*top*), and an uncharacterized, inefficiently-spliced lincRNA, *LINC01029* (*bottom*), in K562 cells. **E.** Mean ESE density per bp in lincRNA and mRNA 3’ (*left*) and 5’ (*right*) splice sites, for all annotated exons larger than 200 bp, compared to random intergenic regions of the same length. **F.** Distribution of ESE density per gene in efficiently (>0.5) and inefficiently spliced (<0.5) lincRNAs.

Strikingly, splicing efficiency was the most discriminatory feature between lincRNAs and mRNAs (Figure 1C). As expected, in the cytosolic fraction, the splicing efficiencies of all RNAs were higher, though lincRNAs were still less efficiently spliced than were mRNAs (Figure 2B and Supplemental Fig S12B). We found no relationship between splicing efficiency and presence of H3K9me3 (FDR<0.05; Supplemental Fig S15). Notably, functionally characterized lincRNAs exhibited greater splicing efficiency than non-characterized lincRNAs with similar expression levels (Figure 2C-D and Supplemental Fig S14). Thus perhaps—similarly to the abundance of conserved TFBSs—splicing efficiency might represent a principal consideration when identifying functional lincRNAs.

To exclude the possibility that the apparent splicing inefficiency derived from inaccuracies in our lincRNA gene models, we repeated our analysis using 3 different gene sets with more reliable annotations. Specifically, we selected: (i) genes annotated as “known” in GENCODE (Supplemental Fig S16), (ii) genes expressed at greater than 1 FPKM (Supplemental Fig S17), and (iii) genes with splicing efficiency larger than 0.1 (Supplemental Fig S18). In 20 out of the 21 comparisons (3 gene sets per 7 cell lines), splicing efficiency was lower in lincRNAs, consistent with our initial observation. Indeed, we also observed – by performing a similar analysis using nuclear and cytosolic RNA-seq data from mouse embryonic stem cells – that murine lincRNAs were less efficiently spliced than control mRNAs (Supplemental Fig S19). This would suggest that inefficient splicing might be a common feature of lincRNAs across species.

Last, we sought to determine if there might be sequence features that could account for the reduction in lincRNA splicing efficiency. To address this we determined if there were differences in exonic splicing enhancers (ESEs), sequence motifs located within exons that regulate alternative splicing (Blencowe 2000). ESEs have been shown to be conserved in lincRNAs (Haerty and Ponting 2015; Schüler et al. 2014). We found that ESE density in lincRNAs was higher than in mRNAs, both in human (Figure 2E) and mouse (Supplemental Fig S20), and that this could be partially explained by differences in GC content (Supplemental Fig S21). Efficiently spliced lincRNAs did not have more ESEs than inefficiently spliced lincRNAs (one-tailed Wilcoxon test P>0.05) (Figure 2F; Supplemental Fig S22) indicating that ESE density cannot account for differences in mRNA vs. lincRNA splicing efficiency.

### The U1-PAS axis is similar in lincRNAs and mRNAs

Another possible factor influencing splicing efficiency could be the binding of the U1 small nuclear ribonucleoprotein (that recognizes and binds the 5’ splice site, and facilitates the recruitment of the spliceosome to the unspliced transcript). Binding of U1 to nascent RNA transcripts is also thought to occlude alternative polyadenylation signals (PAS), thereby protecting the transcript from degradation and discriminating *bona fide* transcripts from transcriptional noise (Almada et al. 2013).

We first evaluated whether presence of canonical U1 motifs was related either to splicing efficiency or to lincRNA locus length. In most cell lines, the presence of a canonical U1 site within a lincRNA—either within the first 1Kb (Figure 3A; Supplemental Fig S23) or within all 5’ splice sites (Spearman’s rho; FDR>0.05)—did not significantly correlate with the efficiency with which that RNA was spliced. However, all RNAs with U1 sites within the first 1Kb were longer than those without, and this difference was greater for lincRNAs than for mRNAs (Figure 3A).

**Figure 3.**
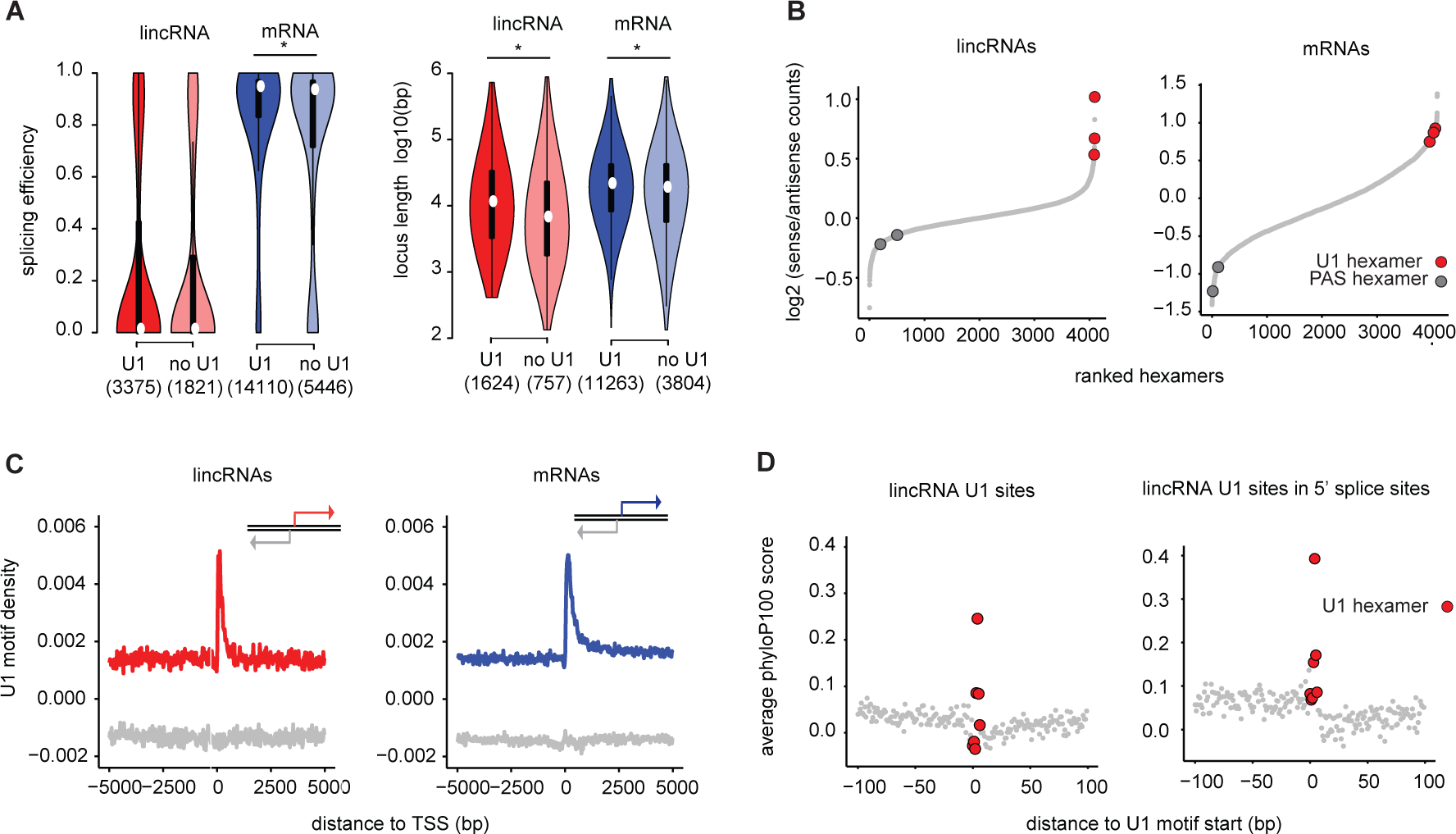
The U1-PAS axis is similar in lincRNAs and mRNAs. **A.** Distribution of splicing efficiency (left) and locus length (right) in lincRNAs and mRNAs in K562, with and without canonical U1 motifs at 5’ splice sites within the first 1Kb downstream of the TSS. **B.** Rank of all hexamers by enrichment in the first 1kb in the sense direction relative to upstream antisense direction in lincRNAs (left) and mRNAs (right). **C.** U1 motif density around the TSS in lincRNAs (left) and mRNAs (right) in the sense strand (red or blue) and in the antisense strand (grey). **D.** Average conservation at all U1 nucleotides present in the first 1 kb of lincRNAs (left) or the subset of these U1 nucleotides that overlap 5’ splice donors (right).

Next, we used a discriminative hexamer analysis (Almada et al. 2013) to assess whether lincRNAs exhibited the so-called “U1-PAS” axis—an enrichment of U1 binding motifs and depletion of polyadenylation motifs downstream of the TSS. In general, the U1-PAS properties of lincRNAs largely mirrored those of their mRNA counterparts. For example, U1 motifs were enriched and polyadenylation motifs were depleted in the sense strand of lincRNA genes, relative to the upstream antisense strand (Figure 3B). Moreover, U1 motifs were highly enriched in the 200 bp downstream of the TSS in both lincRNAs and mRNAs (Figure 3C, Supplemental Table S9). Finally, as in mRNAs, lincRNA U1 motifs were more conserved than were nearby sequences; these U1 conservation signals were further strengthened when located at annotated exon-intron junctions (Figure 3D). We thereby conclude that features of the U1-PAS axis cannot account for inefficient lincRNA splicing.

### Internal splicing signals are weaker in lincRNAs than in mRNAs

Thus far, we have not been able to attribute differences in splicing efficiency between lincRNAs and mRNAs to differences in ESE density, the U1-PAS axis, or the canonical U1 5’ splice site. We therefore investigated whether differences in other splicing signals, such as the polypyrimidine tract (PPT) and the branch point (Figure 4A), might explain this phenomenon. The PPT signals of lincRNAs had a slightly smaller proportion of pyrimidines (Figure 4B; Supplemental Fig S24) and a larger ratio of uracils to cytosines (Figure 4C, D) than those of their mRNA counterparts. In most cell lines, considering both mRNAs and lincRNAs, the overall number of pyrimidines positively correlated with splicing efficiency (Supplemental Fig S25). Moreover, constitutive splice sites exhibited a larger proportion of pyrimidine nucleotides than did those of alternatively spliced sites in both lincRNAs and mRNAs (Figure 4E).

**Figure 4.**
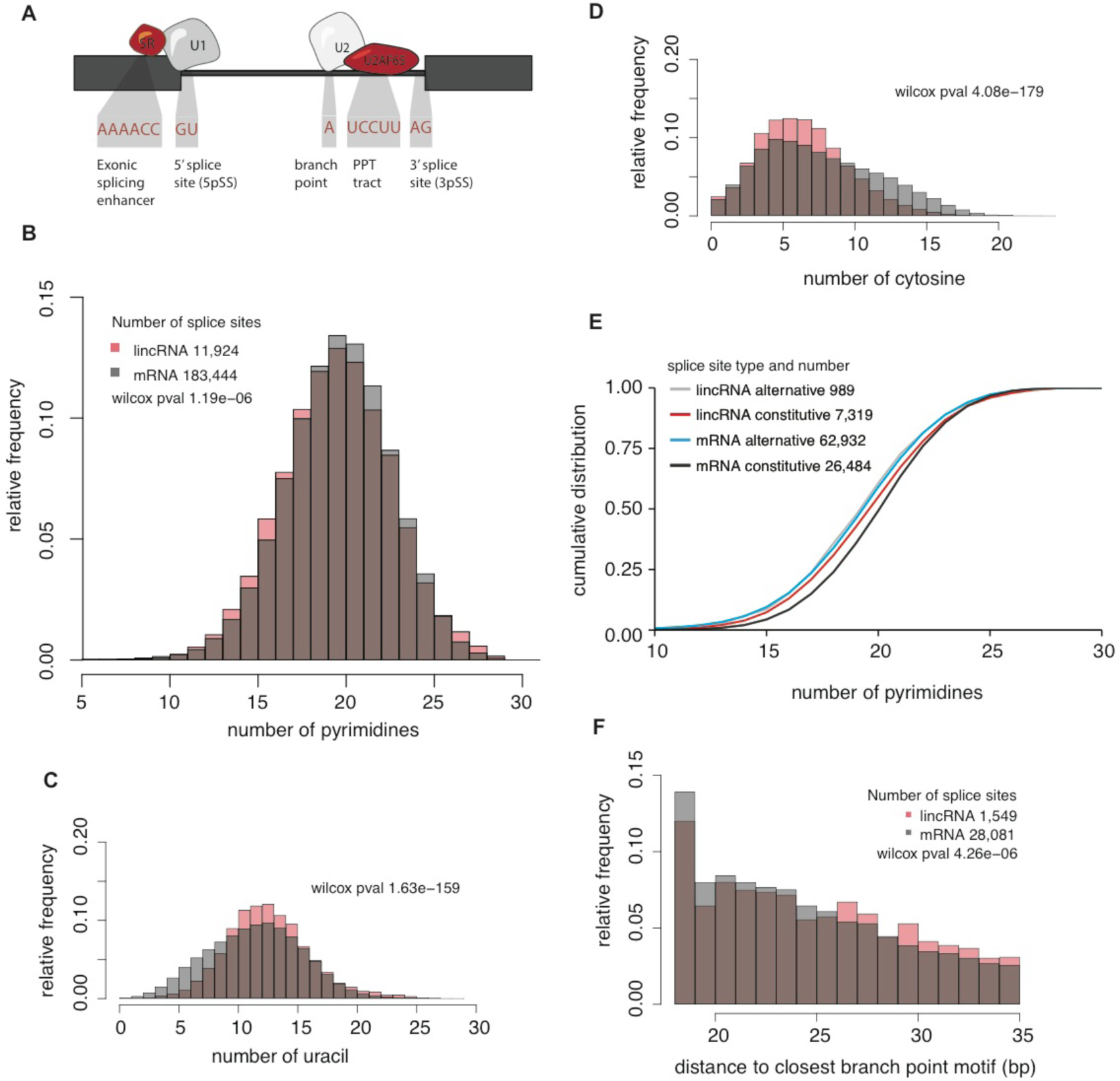
Internal splicing motifs are slightly weaker in lincRNAs than in mRNAs. **A.** Schematic view of necessary splicing RNA motifs and splicing regulators. **B.** Relative frequency of pyrimidine nucleotides upstream of the 3’ splice sites. **C.-D.** Relative frequencies of uracil (C) and cytosine (D) nucleotides upstream of the 3’ splice site. Number of 3’ splice sites analyzed is the same as in B. **E.** Relative frequency of pyrimidine nucleotides upstream of the 3’ splice site grouped by splice site type (alternative or constitutive). **F.** Distribution of distances (bp) between 3’ splice site and the nearest canonical branch point motif.

We next investigated the distance between the branch point and 3’ splice sites, which has been previously shown to affect splicing efficiency (Rosenberg et al. 2015). These distances were greater in lincRNAs than in mRNAs, as gauged both by *in silico* mapping the canonical branch motif (Figure 4F; Supplemental Fig S26) and as observed in CaptureSeq data targeting splicing branch points (Mercer et al. 2015) (Supplemental Fig S27). In summary, number of pyrimidines and branch point differences within internal 3’ splice sites could account for some of the splicing differences between lincRNAs and mRNAs.

### Binding of U2AF65 and splice site conservation correlates with greater splicing efficiency in lincRNAs

Based on the above observations, we hypothesized that weak internal 3’ splice-site signals may contribute to the lower splicing efficiency of lincRNAs. One important factor in splice site determination is U2AF65, which binds to the PPT and promotes binding of the U2 snRNP to the branch point (Figure 4A). Consistent with the known role of U2AF65, we observed that U2AF65 peaks were enriched near 3’ splice sites in both lincRNAs and mRNAs in two publicly available U2AF65 CLIP–seq datasets (Zarnack et al. 2013; Shao et al. 2014) (Figure 5A, B). While U2AF65 binding was depleted in lincRNAs relative to mRNAs in both datasets (Figure 5 A, B), lincRNAs and mRNAs with similar expression levels exhibited similar peak densities (Supplemental Fig S28).

**Figure 5.**
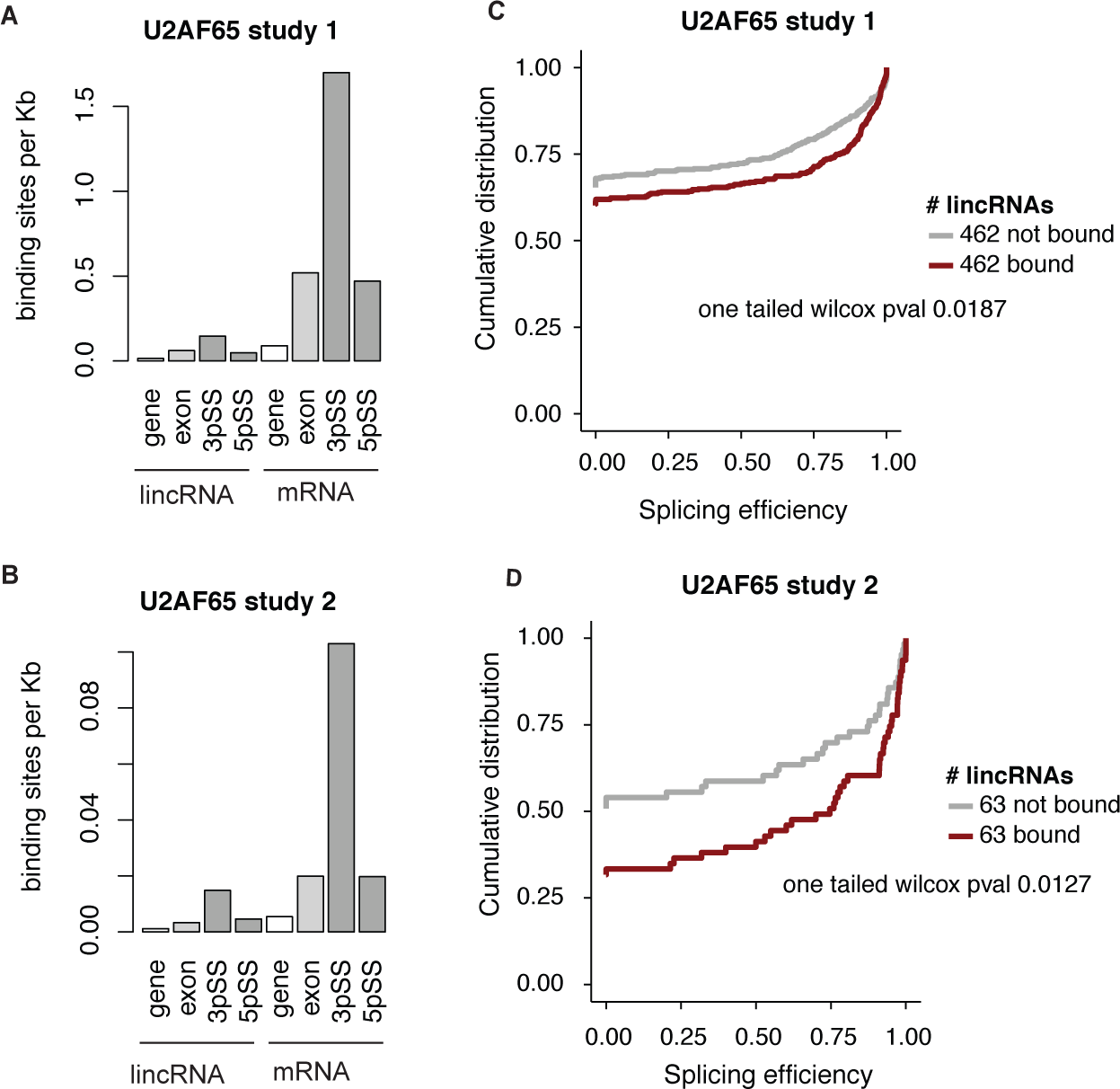
U2AF65 binding in lincRNAs and mRNAs. **A.-B**. U2AF65 CLIP-Seq binding site density in lincRNAs (*left*) and mRNAs (*right*), quantified across the entire gene locus, the processed transcript (exons), and the 3’ and 5’ splice sites. Splice sites were defined as +/−50 bp from any splice junction. **C.-D.** Cumulative distribution of splicing efficiency comparing U2AF65-bound or expression-matched unbound lincRNAs. Study 1 corresponds to data from Zarnack et al. 2013 and study 2 to Shao et al. 2014.

To address whether U2AF65 binding was related to splicing efficiency, we compared the splicing efficiencies of expression-matched lincRNAs that were or were not bound by U2AF65. In both CLIP datasets, U2AF65-bound lincRNAs exhibited significantly greater splicing efficiencies than their unbound counterparts (one tailed Wilcoxon P<0.05, Figure 5 C, D). We observed similar results for lincRNAs with one 3’ splice site (Supplemental Fig S29A), or when lincRNAs with zero splicing efficiency were excluded from analysis (Supplemental Fig S29B). Likewise, U2AF65-bound mRNAs were also more efficiently spliced than unbound mRNAs (one tailed Wilcoxon P<0.05; Supplemental Fig S30). Thus, less U2AF65 binding may contribute to poor splicing in lincRNAs.

We next examined the relationship between lincRNA splicing efficiency and the conservation of 5’ and 3’ splice sites (Supplemental Fig S31). On average, both 5’ and 3’ splice junctions were significantly conserved in lincRNAs (Empirical P<0.011 and P<0.005 respectively). Moreover, 1,105 lincRNAs (21%) had at least one significantly conserved (5’ or 3’) splice site (Supplemental Table S9). These lincRNAs were more efficiently spliced than those lacking conserved sites, even when correcting for expression levels (Supplemental Fig S32). Furthermore, the set of lincRNAs with conserved splice sites (5’ or 3’) was enriched for functionally characterized lincRNAs (Fisher’s exact test P=1.7e-05 and P=0.034 respectively), indicating that splice site conservation may provide a further metric by which to computationally identify functional lincRNAs.

### LincRNAs and mRNAs exhibit similar stability

To complete our systematic survey of the properties associated with the lincRNA lifecycle, we next focused on RNA decay. LincRNAs have been previously observed to be less stable than mRNAs (Clark et al. 2012), although an analysis that incorporates expression levels into assessments of stability has not yet been performed. To address this, we systematically measured the half-lives of lincRNAs and mRNAs following actinomycin D treatment (methods) in two human cell lines: K562 and human embryonic stem cells (HUES9). Strikingly, when we normalized for pre-treatment expression levels, we found that the half-lives of lincRNAs were indistinguishable from those of mRNAs in both cell lines (one tailed Wilcoxon P>0.05) (Figure 6A, Supplemental Fig S33, Supplemental Table S10).

**Figure 6.**
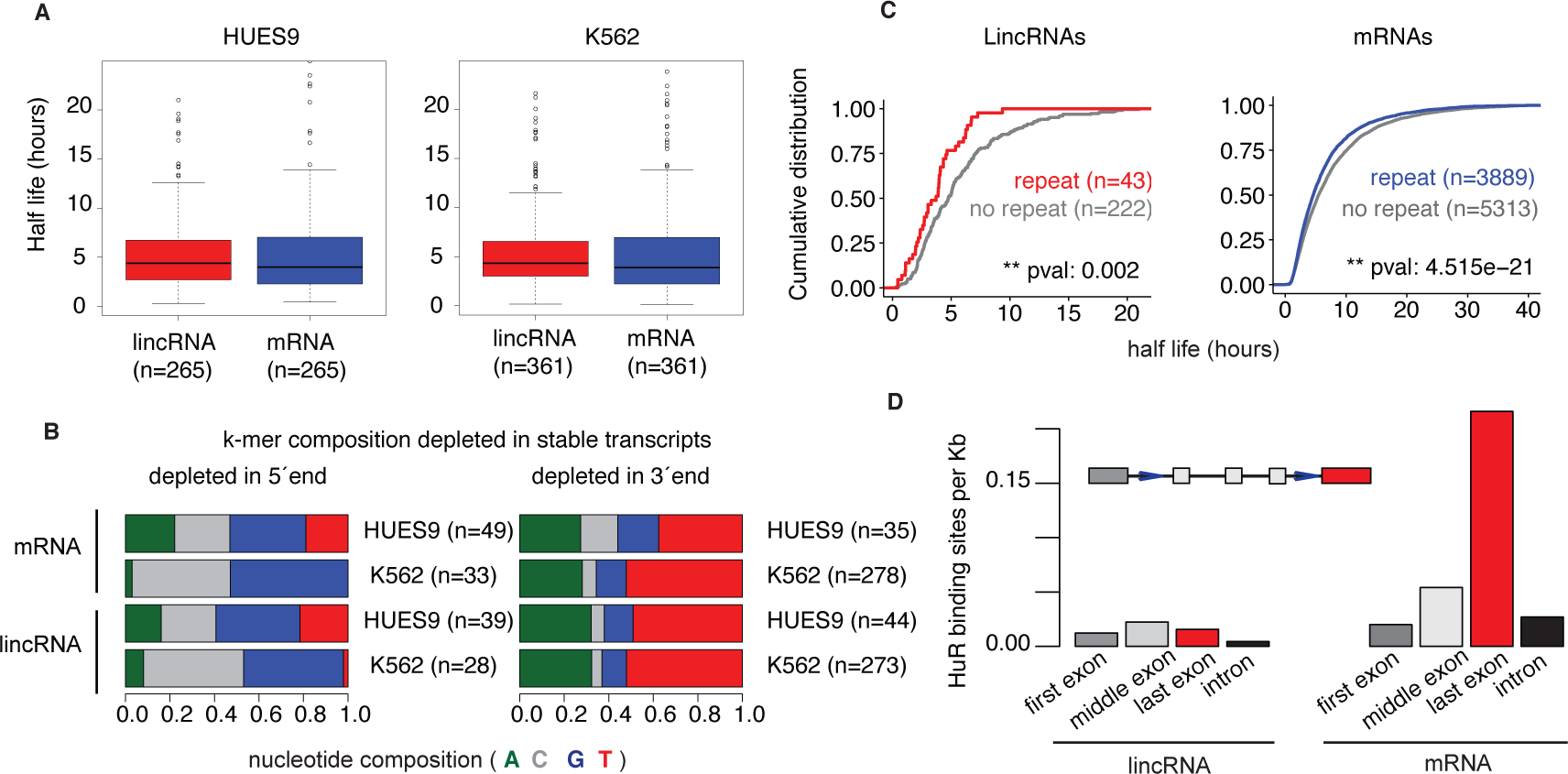
lincRNAs are as stable as expression-matched mRNAs. **A.** Half lives of expression-matched lincRNAs (red) and mRNAs (blue) after actinomycin treatment in HUES9 (*left*) and K562 (*right*) cells. **B.** Nucleotide composition of *7*-mers that are depleted in stable mRNAs and lincRNAs in both the 5’ (left) or 3’ ends (right). *7*-mers were called as significant at FDR<0.05 for mRNAs and at p-values<0.05 for lincRNAs, due to their low sample size. **C.** Cumulative distribution of half lives for lincRNAs (*left*) and mRNAs (*right*) in HUES9 with and without at least one exon that intersects low complexity repeats **D.** Density of HuR binding sites in different regions of lincRNA (left) and and mRNA (right) transcripts.

Furthermore, sequence elements that correlated with RNA stability were similar among lincRNAs and expression-matched mRNAs. We examined the nucleotide composition of *7*-mers that were significantly different between stable and unstable transcripts. Sequences depleted at the 5’ end of all stable transcripts (lincRNAs and mRNAs) were CG rich, while those at the 3’ end were AU rich, in both cell lines (Figure 6B; Supplemental Fig S34). However, the specific *k*-mers that were enriched in lincRNAs and mRNAs differed (Supplemental Fig S35, S36). In addition, in HUES9 cells, low complexity repeats were enriched in both unstable lincRNAs and mRNAs, while in K562 cells this enrichment was only observed in unstable mRNAs (Figure 6C; Wilcoxon test P=0.002; Supplemental Fig S37, S38).

Finally, we assessed how HuR, a known stability regulator that binds to RNA transcripts, may influence lincRNA half-life, using CLIP-seq data (Kishore et al. 2011). We found that, while HuR preferentially bound near the 3’ ends of mRNAs, such positional bias did not exist in lincRNAs regardless of whether the comparison was for expression-matched transcripts (Supplemental Fig S39) or not (Figure 6D, Supplemental Fig S40).

## DISCUSSION

Here we have provided a comprehensive characterization of the pre‐, co‐ and post-transcriptional regulation properties of lincRNAs relative to those of mRNAs. By systematically surveying key aspects of the lincRNA lifecycle—from synthesis to degradation—and comparing them to equivalent stages of the mRNA lifecycle, we have elucidated many features that distinguish these two classes of RNA (Figure 7). In addition, we have identified characteristic signatures that are enriched in known functional lincRNAs.

**Figure 7.**
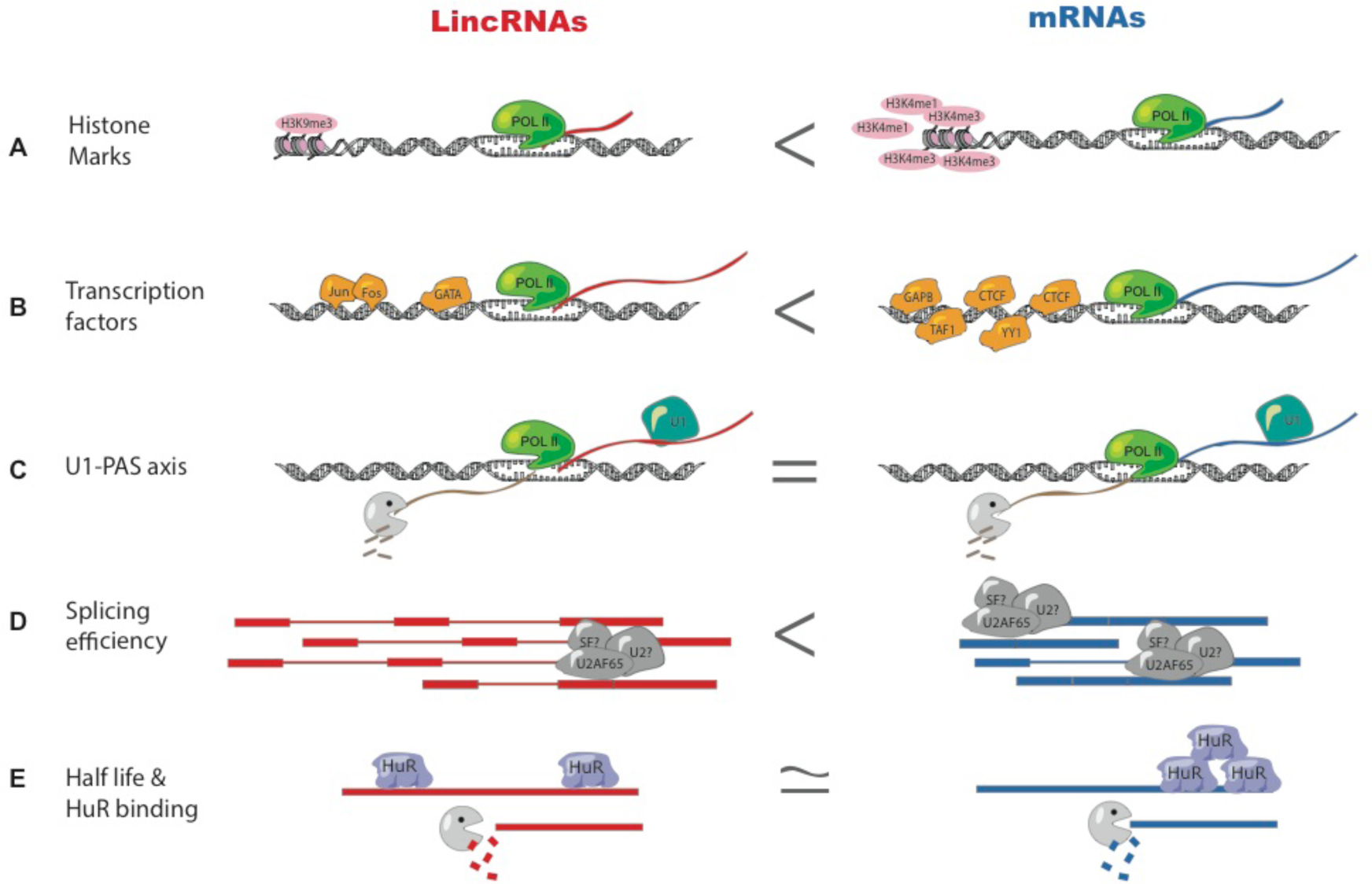
Summary of the similarities and differences between the lincRNA and mRNA life cycles. **A.-B.** LincRNAs have fewer histone marks (A) and transcription factors (B) bound in their promoter than mRNAs. **C.** U1 motif profiles are similar in lincRNAs and mRNAs. **D.** Splicing efficiency is lower in lincRNAs than in mRNAs and lincRNAs are depleted for U2AF65-binding. **E.** LincRNA and mRNA stability is similar when comparing expression-matched groups. HuR equally binds equally to both classes but is biased towards 3’ UTR localization in mRNAs alone.

The first set of properties that distinguish lincRNAs from mRNAs relates to histone modifications and promoter regulation. Previous reports have suggested that transcribed lincRNAs and mRNAs had similar histone marks (Guttman et al. 2009). Whereas in general terms lincRNAs and mRNAs share similar histone marks such as H3K4me3 in their promoter and H3K36 in the gene body, when comparing expression-matched populations, there are slight differences. Possibly the most striking difference is that that H3K9me3—a canonically repressive histone mark—is enriched in the promoters of active lincRNAs (Alam et al. 2014) compared to active mRNAs, and that expression of H3K9me3-marked lincRNAs is more tissue specific than that of lincRNAs lacking H3K9me3. In addition, lincRNAs have fewer transcription factors bound to their promoters than similarly expressed mRNAs. Since our analysis controlled for RNA expression levels, these observations are likely to reflect fundamental differences in the regulation of lincRNA promoters. Whereas mRNAs may require relatively high numbers of TFs to maintain baseline levels of expression across many tissues, lincRNA promoters may generally be in a more repressed state, and are only activated in certain tissues or at certain developmental time points.

Earlier studies suggested that lincRNA promoters are more conserved than mRNA promoters (Carninci et al. 2005), although later work has reported the opposite (Kutter et al. 2012). Here, we find that lincRNA promoters contain many conserved transcription factor binding sites (TFBSs). In fact, the majority of lincRNA loci have at least one conserved TFBS, suggesting that TF binding at lincRNA promoters is functionally relevant. Furthermore, in some instances these TFBSs are more conserved in lincRNAs than in mRNAs. For example, GATA2, one of the main regulators of hematopoiesis, is more enriched and conserved in lincRNA promoters than in mRNA promoters. Additionally, the number of conserved TFBSs is greater in functionally characterized lincRNAs, suggesting that regulation at the promoter level should be taken into account when selecting candidate lincRNAs for follow-up studies.

The most distinctive feature of lincRNA biogenesis pertains to the efficiency and regulation of splicing (Figure 1C). Previous studies suggested that lincRNAs were inefficiently spliced (Tilgner et al. 2012), but had not investigated potential causes of this inefficiency. We have found that lincRNAs have slightly weaker splicing determinants, such as branch point position and PPT sequence composition, and lower binding of the splicing factor U2AF65. It will be intriguing to see if this observation is paralleled in the binding profiles of other core or auxiliary splicing factors. Moreover, this suggests that splicing may not be required for a substantial proportion of lincRNAs and that the annotations of many lincRNAs might require additional unspliced or partially-spliced isoforms. One of the lincRNAs that was first identified as being inefficiently spliced was *Airn*, for which the act of transcription, rather than the RNA molecule itself, has since been shown to be the biologically functional unit (Latos et al. 2012).

However, we also identified a substantial proportion of lincRNAs that are efficiently spliced with conserved splice junctions. Interestingly, efficient splicing is more prevalent among lincRNAs that have been shown to have specific functions, such as *XIST* (Cerase et al. 2015), *FIRRE* (Hacisuleyman et al. 2014), and *MIAT* (Liao et al. 2016). Indeed, efficient splicing has been shown to be necessary for some lincRNAs to carry out their function (Marquardt et al. 2014). Therefore, our results suggest that efficient splicing is an important step in the processing of a subset of lincRNAs, perhaps those that play important roles as RNAs within the cell.

While our analysis has revealed many distinctive characteristics of lincRNA biogenesis, other aspects of post-transcriptional processing have been demonstrated to be similar between lincRNAs and mRNAs. First, canonical U1 sites are as common in lincRNAs as in mRNAs. This finding differs from the observation that lncRNAs arising from divergent mRNA promoters have fewer U1 sites than their mRNA counterparts (Almada et al. 2013) and suggests that lincRNAs and divergent lncRNAs have different properties. Of interest, lincRNAs with U1 sites are longer than those without such sites, implying that U1 binding is likely to protect nascent lincRNA transcripts from premature degradation, as has been shown for mRNAs (Almada et al. 2013). Furthermore, we find that stabilities of lincRNAs resemble those of expression-matched mRNAs. The only feature discriminating these two classes of molecules pertained to the position—and not the frequency—of stability regulator binding.

Our data support a model wherein the degree of lincRNA post-transcriptional regulation is extremely variable. On one end of the distribution, we find lowly expressed, inefficiently spliced lincRNAs with relatively few conserved promoter TFBSs. Despite having RNA regulatory motifs such as U1, we find no evidence of these lincRNA transcripts being under evolutionary constraint. On the other end, we find highly regulated lincRNAs with efficient splicing, conserved exon-intron junctions, and promoters with several conserved TFBSs. Whereas the former could be the result of transcriptional noise or the byproduct of a necessary act of transcription (Melé et al. 2016), the latter group of more regulated lincRNAs are better candidates to function as RNA molecules. Here, we provide a list of these tightly regulated lincRNAs for future analysis and validation (Supplemental Table S11).

Collectively, these observations reveal important insights into how lincRNA biogenesis is regulated and provide several possible explanations for the differences between lincRNAs and mRNAs—all of which can be tested experimentally. Moreover, this work highlights the substantial variability of transcriptional regulation across lincRNAs and provides novel criteria with which to select candidate functional lincRNAs for future validation.

## METHODS

### LincRNA and mRNA selection in human and mouse

We analyzed all genes in the human GENCODE v19 catalog with a biotype of “lincRNA” or “protein_coding” (Harrow et al. 2012). We excluded lincRNAs that were located within 5kb of any protein coding gene, and any protein coding genes whose promoter (+/−5kb of TSS) overlapped a lincRNA promoter. We also excluded any annotated lincRNAs with protein-coding potential greater than zero according to PhyloCSF (Lin et al. 2011). Transcript-level analyses were performed on each gene’s longest transcript. For mouse, we analyzed all genes in the GENCODE vM6 with a biotype of “lincRNA” or “protein_coding”. We ignored all lincRNAs with protein coding probability greater than 0.44 according to CPAT (Wang et al. 2013). To select a set of lincRNAs with validated functions, we selected those lincRNAs with an Ensemble gene_id in lncRNAdb (Amaral et al. 2011) (www.lncRNAdb.com).

### ENCODE expression quantification

To quantify gene expression, we downloaded publicly available RNA-seq data from 11 ENCODE cell lines (GSE30567; polyA+, whole-cell, nucleus and cytosol samples: A549, GM12878, H1-hESC, HeLa-S3, HepG2, HUVEC, IMR90, K562, MCF-7, NHEK, SK-N-SH). We mapped reads to the human genome (hg19) using TopHat v2.1.0 (Trapnell et al. 2009; Kim et al. 2013) with the “‐‐no-coverage-search” flag and using GENCODE v19 annotation. We quantified expression with Cuffdiff2 (Trapnell et al. 2012). We took a gene or transcript’s expression level as the average FPKM level across replicates. We calculated nuclear/cytosolic ratio by dividing the gene expression levels (as fragments per kilobase per million reads, FPKM) in each cell compartment. We considered any gene to be expressed in a particular cell line if its expression levels were > 0.1 FPKM, similar to what has been used in other large-scale transcritpomics studies (Melé et al. 2015).

### Tissue specificity

We used 16 tissues from the Illumina Human Body Map 2.0 (GSE30611; Adipose, Adrenal, Brain, Breast, Colon, Heart, Kidney, Liver, Lung, Lymph Node, Ovary, Prostate, Skeletal Muscle, Testes, Thyroid, White Blood Cells) and four tissues from Cabili et al (2011) (GSE30554; Foreskin Fibroblast, HeLa, Lung Fibroblast, Placenta). We mapped and quantified the raw reads as described above. We then defined an entropy-based “tissue specificity score” for each gene as described previously (Cabili et al. 2011).

### ChIP-seq analysis

We used publicly available ChIP-seq data from ENCODE (www.encodeproject.org). We downloaded broadPeak and narrowPeak files for 91 cell lines from UCSC (Supplemental Table S12) and selected the first replicate from each experiment. We defined promoters as the regions −/+ 5000 or −2000/+1000 bp of a TSS. For each TF or histone modification, we used BedTools (Quinlan and Hall 2010) to intersect the ChIP peaks with the promoters and summed up the number of peaks overlapping each gene’s promoter.

### Splicing efficiency

We estimated transcript and gene abundances using a modified annotation that contained an additional isoform per gene spanning the gene locus. We calculated splicing efficiency as the sum of abundances of all annotated isoforms divided by the sum of abundances of all isoforms including the spanning one (Supplemental Fig S11A). We removed single-exon genes from the analysis. For analysis relying on splice site annotation and to avoid spurious results due to misannotations, we replicated our analyses excluding those lincRNAs that had splicing efficiency equal to zero in cell lines for which splicing efficiency could be calculated.

### Nuclear fractionation of mouse ES cells

We isolated cytoplasmic and nuclear fractions from N2 mESC cells as described (Hacisuleyman et al. 2014) for mESC cells. We prepared poly(A)+ mRNA-seq libraries using the TruSeq RNA sample preparation kit, v2 (Illumina) as described (Goff et al. 2015). Details of the protocol can be found in Supplementary Methods.

### Effect size calculation

For continuous values, we used Wilcoxon’s effect size formula:

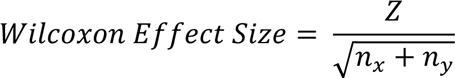

where *Z* is the test statistic and *n* is the number of observations in the *x* and *y* groups that we are comparing. To assess effect size on binary properties such as presence/absence of specific transcription factors or histone marks bound, we used the phi coefficient from the psych package (function: phi) (R Core Team 2014).

### Expression matching

We used the R package matchIt with default settings (Ho et al. 2011) where, for each lincRNA, the mRNA with the closest expression value is selected.

### Conservation analyses

We used the vertebrate 100-way alignment PhyloP scores (Pollard et al. 2010) and extracted the scores from genomic regions using bwtool (Pohl and Beato 2014).

#### TF conservation

For each TF, we centered the ChIP-seq peaks overlapping a lincRNA or mRNA promoter at the peak maxima and merged any peaks closer than 50 bp. We then compared the 200 bp region centered on the peak maxima to the 100 bp regions at each side and at a distance of 150 bp from the peak as background. To test conservation at single ChIP-seq peaks, we performed the same test for each peak individually. To compare ChIP-seq peaks overlapping a known TF motif, we mapped known motifs using FIMO (Grant et al. 2011) and intersected these with corresponding ChIP peaks using bedtools.

#### U1 sites, 3’ and 5’ splice site conservation

We compared the average score of canonical U1 sites, 3’ and 5’ splice site *k*-mers to all other adjacent *k*-mers in a 200 nt window. We considered a *k*-mer to be conserved if it had a greater average conservation score than 95% of all other adjacent *k*-mers. We also performed the same test for each sequence individually. We used *2*-mers for 3’ splice sites, and *10*-mers for 5’ splice sites to account for all potential U1 hexamers in splice donor sites.

### Motif analyses

#### TF binding motifs

We downloaded all TF binding motifs from the JASPAR CORE database (Mathelier et al. 2016). We mapped them to the corresponding ChIP-seq peaks that overlapped lincRNA or mRNA promoters using the FIMO program (Grant et al. 2011) from the MEME package (Bailey et al. 2009).

#### ESEs

We downloaded ESE motifs for both humans and mice from (Fairbrother et al. 2002). To calculate ESE density, we used all annotated unique exon-intron junctions in both lincRNAs and mRNAs, where the exons were at least 200 nt long.

#### U1/PAS

We used three canonical U1 sites (GGUAAG, GGUGAG, GUGAGU) and two PAS motifs (AAUAAA, AUUAAA) as in Almada et al. (2013).

#### Polypyrimidine tract

We counted the number of pyrimidine nucleotides (cytosine or uracil) in a region of 30 nucleotides upstream from all annotated 3’ splice sites. **Branch point**: We mapped the canonical branch point sequence (CU[AG]A[CU]) to the region between every annotated 3’ splice site and 40 bp upstream.

#### 3 prime splice sites

We selected all annotated non-redundant 3’ splice sites in lincRNAs and mRNAs. We defined constitutive or alternative 3’ splice sites as those present in at least 75% or in less than 25% of all annotated isoforms of a gene respectively.

### *K*-mer analyses

To perform discriminative *k*-mer enrichment analyses, we used the program Jellyfish (Marçais and Kingsford 2011). For each *k*-mer, we then calculated the log2 fold ratio between sequences of interest: either *k*-mers enriched in the downstream sense direction of a TSS as compared to the upstream antisense direction or *k*-mers enriched in stable versus unstable transcripts (details in supplementary methods)

### RNA stability assay

We treated K562 and human embryonic stem cells (HUES9) with actinomycin D (5 μg/ml of culture medium) and harvested the cells at 0 min, 30 min, 2h, 4h and 8h for RNA isolation and measured global RNA levels by poly(A)+ RNA-seq in triplicate. For each gene, we normalized abundance to GAPDH and fitted the data to a first order exponential decay curve. We then calculated the average half-life per gene across replicates in which the Pearson correlation between the fitted curve and the real curve was higher than 0.7.

### CLIP-seq data analysis

We downloaded CLiP-seq from publicly available data for U2AF65 from HeLa cells from two datasets: (Zarnack et al. 2013) (ArrayExpress accession numbers for the iCLIP data: E-MTAB-1371) and (Shao et al. 2014) (Gene Expression Omnibus GSE61603). We used an established bioinformatics pipeline designed to find CLIP-seq peaks relative to the transcript abundances (Kelley et al. 2014) and calculated peak density across different transcript regions (further details in supplementary methods).

## DATA ACCESS

The sequencing data from this study have been submitted to the NCBI Gene Expression Omnibus (GEO; http://www.ncbi.nlm.nih.gov/geo/) under accession number GSE80046.

## ACKNOWLEDGMENTS

We thank David Kelley from the Rinn lab for feedback on CLIP-seq analysis and comments on the manuscript, James Lee for comments on the manuscript, Alex Meissner for providing the HUES9 cells and Veronica Akopian for technical support with HUES9 culture.

Marta Melé Messeguer is a Gilead Fellow of the Life Sciences Research Foundation. This work was supported by US National Institutes of Health grants R01 ES020260, R01 MH102416, and P01 GM099117 and by the National Science Foundation Graduate Research Fellowship under Grant No. DGE1144152.

## AUTHOR CONTRIBUTIONS

MM and JR designed the project and wrote the manuscript. KM performed all *k*-mer and conservation analysis. WM processed ENCODE RNA-seq and ChIP-seq datasets. DS performed fractionation protocol for mouse ES cells. CG performed the stability assay. MM performed all the remaining analysis. All authors have read and approved the manuscript for publication.

